# Constitutive neural networks for main pulmonary arteries: Discovering the undiscovered

**DOI:** 10.1101/2024.10.31.621391

**Authors:** Thibault Vervenne, Mathias Peirlinck, Nele Famaey, Ellen Kuhl

## Abstract

Accurate modeling of cardiovascular tissues is crucial for understanding and predicting their behavior in various physiological and pathological conditions. In this study, we specifically focus on the pulmonary artery in the context of the Ross procedure, using neural networks to discover the most suitable material model. The Ross procedure is a complex cardiac surgery where the patient’s own pulmonary valve is used to replace the diseased aortic valve. Ensuring the successful long-term outcomes of this intervention requires a detailed understanding of the mechanical properties of pulmonary tissue. Constitutive artificial neural networks offer a novel approach to capture such complex stress-strain relationships. Here we design and train different constitutive neural networks to characterize the hyperelastic, anisotropic behavior of the main pulmonary artery. Informed by experimental biaxial testing data under various axial-circumferential loading ratios, these networks automatically discover the inherent material behavior, without the limitations of predefined mathematical models. We regularize the model discovery using cross-sample feature selection and explore its sensitivity to the collagen fiber distribution. Strikingly, we uniformly discover an isotropic exponential first-invariant term and an anisotropic quadratic fifth-invariant term. We show that constitutive models with both these terms can reliably predict arterial responses under diverse loading conditions. Our results provide crucial improvements in experimental data agreement, and enhance our understanding into the biomechanical properties of pulmonary tissue. The model outcomes can be used in a variety of computational frameworks of autograft adaptation, ultimately improving the surgical outcomes after the Ross procedure.

## 1 Introduction

Automated model discovery with constitutive artificial neural networks (CANNs) is an emerging technology aimed at improving the modeling of complex biological tissues (Linka and Kuhl, 2023; St. Pierre et al, 2024; Martonova et al, 2024). This method combines machine learning with traditional constitutive modeling techniques to automatically discover material laws that describe how biological tissues deform under various conditions. Cardiovascular tissues, such as arteries and heart valves, are highly complex, exhibiting nonlinear and anisotropic behaviors. A priori selection of an appropriate traditional constitutive model to describe these behaviors requires a deep expert knowledge, which can introduce bias and limits successful modeling to a few well-trained individuals. In contrast, CANNs leverage data from experiments or simulations to learn these relationships automatically. By training on experimental data, these networks are capable of capturing the intrinsic stress-strain relationships of cardiovascular tissues and, when done right, even generalize well to predict responses under previously unseen conditions.

Previous research has investigated anisotropic neural network models for human skin and healthy aortic arteries (Linka et al, 2023; Peirlinck et al, 2024b). Informed by biaxial extension tests, these networks inherently satisfied general kinematic, thermodynamic, and physical constraints. The results suggest that constitutive neural networks can robustly and consistently discover variations of arterial models from experimental data. Notably, the discovered models are generalizerable and interpretable, with the majority of network weights training to zero, while a small subset of non-zero weights defines the discovered model (Linka and Kuhl, 2024).

CANNs offer a key advantage in cardiovascular applications by automatically handling the complex, nonlinear nature of tissue mechanics without relying on oversimplified assumptions (Linka et al, 2021, 2022). This holds promise for personalized medicine, where models can be customized to patient data to individualize and improve surgical planning and device design (Peirlinck et al, 2021). The automated nature of this approach reduces the time and expertise needed for developing accurate models. CANNs are designed with specific architectural features to ensure adherence to physics-based constraints inherent to constitutive modeling. Classical artificial neural networks rely heavily on carefully chosen activation functions and architectures that align with the problem at hand. CANNs therefore represent an innovative class of artificial neural networks that are explicitly tailored to satisfy kinematical, thermodynamical, and physical principles by design. They also restrict the space of permissible functions, enabling robust training on experimental data.

In the realm of automated discovery of constitutive models from biological testing data, CANNs utilize specialized inputs (such as deformation gradients) and outputs (such as stress measures derived from an energy density function). Additionally, their activation functions are crafted to uphold essential properties like thermodynamic consistency, material objectivity, material symmetry, physical constraints, and polyconvexity (Linka and Kuhl, 2023).

Therefore, ongoing research can benefit from the tight integration of biomechanics, biomedical engineering, and physics-informed machine learning (Holzapfel and Ogden, 2017). Recent findings indeed suggest that neural networks could automate model discovery and parametrization, potentially transforming constitutive modeling and subsequent finite element simulations (Weiss et al, 1996; Peirlinck et al, 2024a,c). Moreover, the highthroughput results of neural networks allow for an in-depth analysis of the biological nature of the considered data, providing additional insights into the microstructural features of the input data.

The present work focuses on automated model discovery and the application of constitutive neural networks to main pulmonary artery experimental data, with the Ross procedure in mind. The Ross procedure is a surgical technique to treat aortic valve disease, particularly in young and active patients. In the 1960s, Dr. Donald Ross designed this intervention by taking the patient’s own pulmonary valve and placing it in the aortic position to create a pulmonary autograft (Charitos et al, 2012; Chauvette et al, 2019; El-Hamamsy et al, 2022; Van Hoof et al, 2022). The new valve is alive and can adapt over time, making the Ross procedure the only aortic valve replacement technique that can restore long-term survival and maintain quality of life. Despite scientific consensus on its significant benefits, the Ross procedure still has a principal failure mode, linked to wall dilatation, which can lead to valve leakage and reoperation. Dilatation occurs due to the inability of the pulmonary autograft to adapt to the sudden increase in loading when exposed to the four- to fivefold increase in aortic pressures.

Here, we contribute to the broader picture of this research by further investigating the mechanical behavior of pulmonary tissue, which forms the baseline for the long-term outcomes of the autograft in the Ross procedure. Multiple computational modeling studies are currently investigating the growth and remodeling outcomes of the Ross procedure, ranging from analytical solutions to patient-specific finite element frameworks (Famaey et al, 2018; Vastmans et al, 2021; Vervenne et al, 2023; Maes et al, 2024). Ultimately, all of these models and results could make the Ross procedure the preferred solution for aortic valve replacement worldwide (Middendorp et al, 2024). At the very core of these frameworks are the constitutive models and parameters of the pulmonary autograft tissue. The sensitivity of the constitutive equations and mechanical properties considered cannot be underestimated, and the presented CANNs aim to increase the overall biofidelity of these computer simulations.

We base our study on biaxial extension data from *n* = 8 pulmonary artery samples under different loading ratios. We perform sample-specific model discovery both with and without regularization. Additionally, we systematically explore constitutive feature selection across all samples. This will allow us to identify a universal constitutive behavior through cross-sample feature selection regularization. To probe the anisotropic tissue response, we perform a fiber angle sensitivity analysis using a sweep of models with fixed fiber angles across the axial-circumferential domain. Finally, we compare our results with the traditional Holzapfel-Gasser-Ogden model, which considers strain-stiffening collagen fibers in an isotropic elastin matrix (Holzapfel et al, 2000). Ultimately, our discovered models and constitutive insights into main pulmonary arteries can inform and support computational studies, particularly in the context of aortic valve replacements through the Ross procedure.

## 2 Materials and Methods

This section will first provide an overview of the unpublished experimental mechanical biaxial extension data used to train the neural networks. We will then outline the constitutive equations, including the deformation gradient and its invariants. These invariants form the basis of the total strain energy density function that constitutes the core of a constitutive artificial neural network. We will design potential terms of this free energy density based on principles of thermodynamic consistency, material objectivity and symmetry, incompressibility, and polyconvexity. Subsequently, we will highlight the specific axial and circumferential first Piola-Kirchhoff stress calculations for planar biaxial extension tests. At the model input, we will also study the effects of varying collagen fiber angles and assess their influence on the discovered models. Finally, we will outline different variations of the neural network loss function used for training.

### 2.1 Experimental dataset

In the context of an animal study on the Ross procedure (Van Hoof et al, 2021), we performed planar biaxial tensile tests on squared samples (10 mm x 10 mm) of the ovine main pulmonary artery, as shown in Fig. 1. The intimal side, or inner surface, of the pulmonary artery was exposed through an axial cut between the anterior cusp (left side of the figure) and the left cusp (right side of the figure) (Thiene and Rizzo, 2023). Two main pulmonary artery samples (A and B), distal to the valve sinuses, were extracted from a total of four ovine pulmonary roots, bringing the total number of tested samples to *n* = 4 · 2 = 8.

**Fig. 1.**
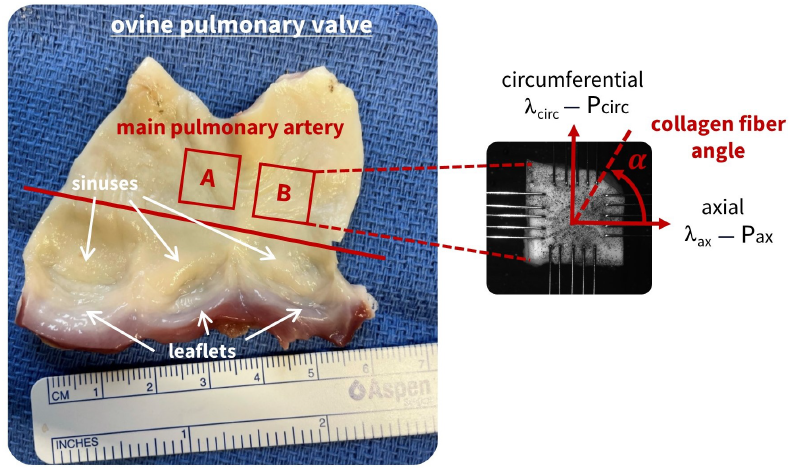
Biaxial tissue extraction and mounting. From four ovine pulmonary roots, two main pulmonary artery squared samples per sheep were extracted. This resulted in a total of *n* = 8 samples, which were mounted in the biaxial test device through rake fixation. The main collagen fiber angle *α* is indicated in the figure and will be varied in the axialcircumferential plane during modeling.

Five rakes were mounted on each side of the sample, with an initial distance of 7 mm between the rakes in both the axial and circumferential directions. Planar biaxial tensile testing (MessPhysik Zwick/Roell) was performed at the Core Facility for Biomechanical Characterization, FIBEr, KU Leuven. Axial and circumferential stretch was increased stepwise until single-rake failure occurred. This procedure resulted in a complete dataset of increasing stretch levels at 5%, 10%, 20%, and 30%.

Native pulmonary arteries operate under diastolic-systolic pressures of 8-20 mmHg, while systemic blood pressures in the aorta average around 80-120 mmHg (Van Hoof et al, 2022).The biaxial load free configuration corresponds to the excised tissue under zero blood pressure, with unknown axial and circumferential prestretches compared to the *in vivo* physiological condition. Our displacement driven test protocol applied 30% deformation to the actuators (*λ*^*^), before preload correction and different from the DIC strain measurements. Debes and Fung (1995) reported 21.5% circumferential strain and 36.5% axial strain for *in vivo* pulmonary arterial tissue, which is in the range of our end-state experimentally biaxial data, with · 50% strains in both directions. This rich dataset is especially important in the context of the Ross procedure, exposing the pulmonary tissue to supra-physiological strains.

Three axial-to-circumferential stretch ratios (2:1, 1:1, 1:2) were applied through the displacement-driven actuators, for each stretch level. A 2:1 ratio means that the full stretch level was applied by the actuator in the axial direction, and half that stretch level in the circumferential direction. Four preconditioning cycles were applied for each loading and stretch ratio, and only the fifth cycle was considered for post-processing (Fehervary et al, 2016).

Here, we write the imposed stretch levels and ratios as applied by the actuators as 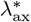 and 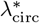,which can be different from the resulting measured planar biaxial deformations in the tissue. Therefore, during testing, 2D digital image correlation (DIC) particle tracking was performed using VIC 2D (Correlated Solutions) to calculate the resulting tissue strains (Vander Linden et al, 2023). We ensure uniform stresses and strains by considering the central 25% of the sample area (Sacks, 2000). The results from opposite actuators were averaged to obtain a single axial and circumferential curve. This resulted in the actual stretches *λ*_ax_ and *λ*_circ_ shown on the x-axes of our plots. Experimentally measured forces were converted to tissue stresses on the y-axes using initial load-free thickness and areal sample measurements, obtained before testing via a Micro Laser Scanner (Acacia Technology).

The resulting engineering stress versus DIC stretch curves for the three ratios and post-preconditioning last loading cycle are presented for one representative sample in Fig. 2. The complete dataset for all *n* = 8 samples is available in the Supplementary Materials. The complete biaxial dataset shows a clear sample-to-sample heterogeneity, even within the same animal. These aleatoric and epistemic uncertainties related to biaxial testing of soft tissues have led to a generalized workflow on all samples, rather than comparing data per sheep. In the axial-circumferential plane, shear stretches were two orders of magnitude lower than the diagonal components of the deformation gradient, highlighting minimal sample alignment error during mounting (Fehervary et al, 2018). The latter is also in agreement with the assumption of symmetrical collagen fiber families around the axial and circumferential directions.

**Fig. 2.**
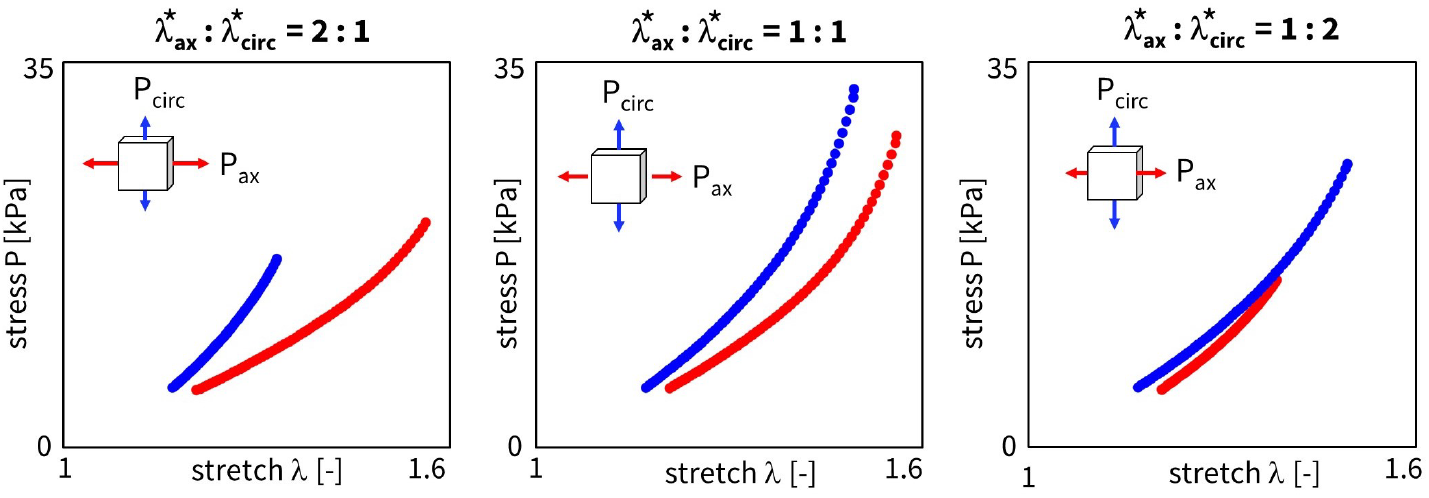
Experimental stress-strain curves. Engineering stress **P** versus diagonal stretch components *λ* from the deformation gradient **F** for a representative sample, in both the axial (red) and circumferential (blue) direction. From left to right, the plots show the axial-to-circumferential ratios of 2:1, 1:1, and 1:2. The curves represent the 5^th^ loading cycle at the 30% stretch level, with the preload correction applied to the stretch domain.

One should note that the engineering stress curves do not start at zero stretch or zero stress. This is due to an applied preload of 0.1 N during testing, which is reasonable for aortic tissues but was considered excessive for pulmonary artery tissues. Therefore, we corrected the zero-strain state and considered the moment in time when all four actuators experienced positive forces as the new reference. The sample-specific starting stress of around 5 kPa in Fig. 2 thus corresponds to the preloaded value 0.1 N, before DIC correction. Note that this force-driven prestretch correction differs in the stretch domain between the axial and circumferential directions, *λ*_ax_ and *λ*_circ_, but uses the same 1:1 ratio offset, 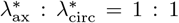, for all three stretch ratios.

### 2.2 Constitutive modeling

In continuum mechanics, we can represent the initial configuration through a Lagrangian variable, ***X***, and the current or deformed configuration by a Eulerian variable, ***x***. A mapping function can transform one state into the other, ***x*** = φ(***X***). Locally, we use the deformation gradient tensor **F** to represent the change in infinitesimal line elements between both configurations, *d****x*** = **F**(***X***)*d****X***. We can then write

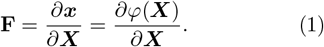

The symmetric right Cauchy-Green deformation tensor **C** is defined as **C** = **F**^*T*^ **F**. We can now indicate the Green-Lagrange strain as 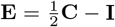, where **I** is the identity tensor (Mooney, 1940; Ciarlet, 1988; Holzapfel, 2002).

Yet another important continuum mechanics variable is the Jacobian *J* = det(**F**). The Jacobian expression itself is in the third invariant, *I*_3_ = det(**F**^*T*^ · **F**) = *J* ^2^. Here, we assume a perfectly incompressible arterial tissue, thus characterized by a constant Jacobian equal to one, *I*_3_ = *J* ^2^ = 1. For an anisotropic material, we can write the isotropic invariants *I*_1_ and *I*_2_ as well as anisotropic invariants *I*_4_ and *I*_5_ as functions of the deformation gradient (Spencer, 1971):

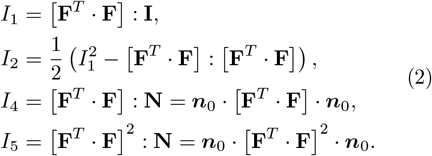

Here, ***n***_0_ is the main fiber direction in the undeformed configuration, and ***n*** is its representation in the deformed configuration. As such, we can write ***n*** = **F** · ***n***_0_, with its associated structure tensor **N** = ***n***_0_ ⊗***n***_0_. Considering a collagen fiber angle *α* within the biaxial plane, we can then state that ***n***_0_ = [cos *α*, sin *α*, 0]^*T*^. This indicative fiber direction is indicated in Fig. 1.

In what follows, and in the context of large deformations, we introduce the first Piola Kirchhoff stress **P** = **P**(**F**) as

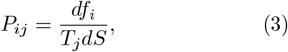

where *df*_*i*_ is the force component in the deformed configuration acting on the corresponding initial surface area. The latter is defined by *dS* as the initial surface area on which the force acts, and with *T*_*j*_ representing the component of the unit normal vector in the *j*-direction in the reference configuration. For readability, we will use the term engineering stress as first Piola Kirchhoff stress alternate. Considering thermodynamic consistency, we imply this engineering stress **P** is equal to the derivative of the strain energy *ψ* with respect to the deformation gradient **F**,

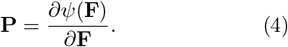

Next, material objectivity enforces that the constitutive equation does not depend on the external frame of reference, and we require that the free energy function *ψ* is a function of the right Cauchy-Green tensor **C** = **F**^*T*^ · **F** (Noll, 1958):

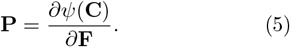

Material symmetry for transversely isotropic materials now allows us to write:

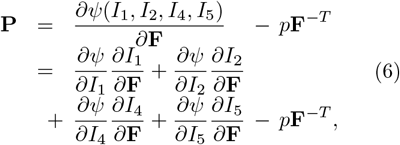

where we again assume perfect incompressibility, enforced by a hydrostatic pressure term − *p***F**^−*T*^. We will be able to solve the latter from boundary conditions later on. Finally, if we consider polyconvexity of the free energy density function by ensuring that *ψ* = *ψ*(*I*_1_) + *ψ*(*I*_2_) + *ψ*(*I*_4_) + *ψ*(*I*_5_), and by deriving the partial derivatives of equations 2 as

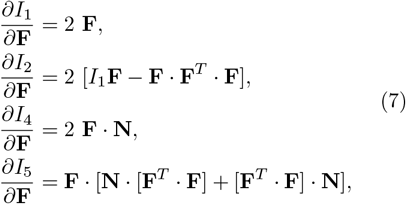

we can write:

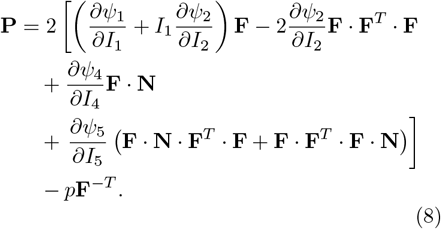

The engineering stress is now defined as a function of the deformation gradient and the derivative of the strain energy density function with respect to the four invariants (Hartmann and Neff, 2003; Ehret and Itskov, 2007).

### 2.3 Constitutive artificial neural network

Fig. 3 shows the constitutive artificial neural network framework with the relevant invariants for transversely isotropic, perfectly incompressible materials. The network has two hidden layers, with four and eight nodes. The first layer generates powers (°) and (°)^2^ of the network input and the second layer applies the identity (°) and exponential functions (exp(°)) to these powers. The network is not fully connected by design to satisfy the condition of polyconvexity a priori. Explicitly, we can now write the resulting strain or free energy density *ψ*(**F**) as a function of the considered invariants:

**Fig. 3.**
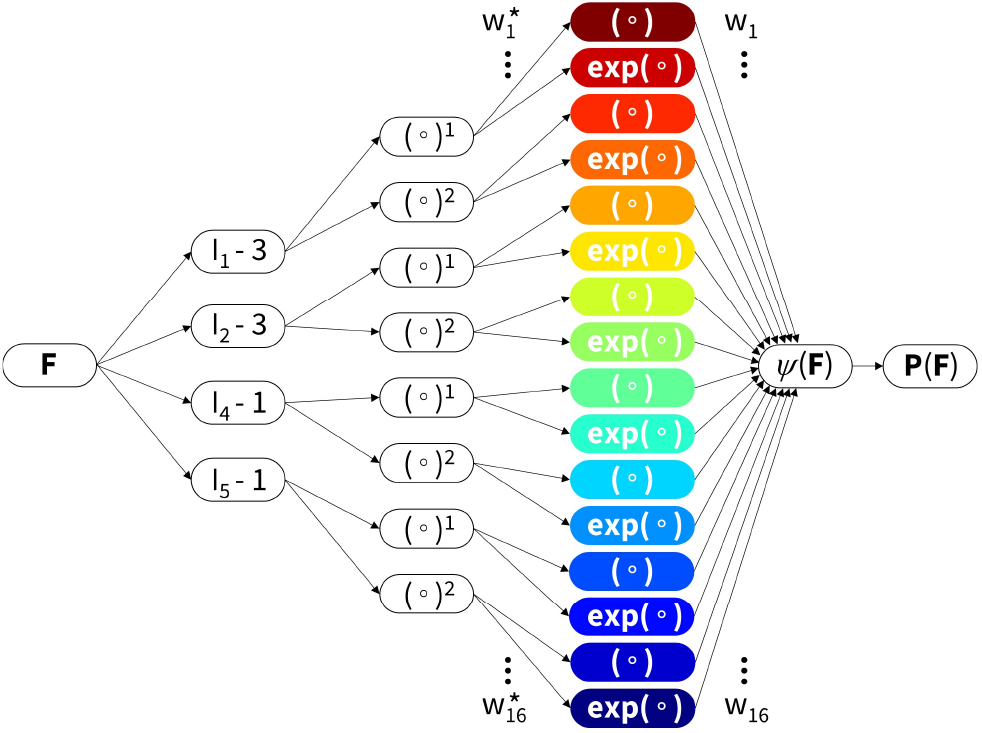
Constitutive Artificial Neural Network. Transversely isotropic, perfectly incompressible, constitutive artificial neural network with two hidden layers for approximating the free energy function *ψ*(*I*_1_, *I*_2_, *I*_4_, *I*_5_) as a function of the invariants of the deformation gradient **F** using sixteen terms. The first layer generates powers (°) and (°)^2^ of the network input and the second layer applies the identity (°) and exponential functions (exp(°)) to these powers. Adopted from (Linka et al, 2023). The chosen color code for the invariants in their specific forms will be respected throughout the continuation of the paper.

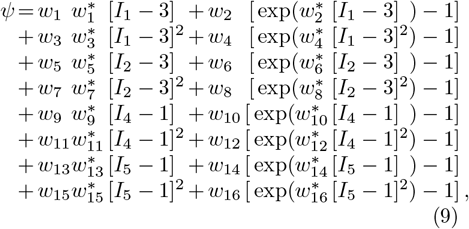

corrected by the pressure term, *ψ* = *ψ* −*p* [*J* −1]. The network has a total of two times 16 trainable weights, with 2^16^ = 65, 536 possible combinations of terms. In other words, we can learn any strain energy function that follows from firstand second order, and identity and exponential additive combinatorics of the *I*_1_, *I*_2_, *I*_4_, and/or *I*_5_ invariants. Note that the growth condition corrections −3 and −1 for the isotropic and anisotropic terms, respectively, ensure that *ψ* = 0 in the undeformed state where **F** = **I**, and thus *I*_1_ = 3, *I*_2_ = 3, *I*_4_ = 1, and *I*_5_ = 1.

Here, we focus on a limited set of polynomial and exponential functions, prescribed prior to model training. Although this limits the model’s learning capabilities, this assumption allows the network weights to have a clear physical interpretation in the form of shear moduli, stiffness-like parameters, and exponential coefficients (Linka and Kuhl, 2023). Therefore, CANNs as new subclass of models still generalize all existing classical constitutive models for arterial tissue.

#### Planar biaxial extension testing

Through experienced mounting of the samples in the test device, as indicated in Fig. 1, we can state that the transversally isotropic pulmonary artery is stretched in two perfectly orthogonal directions, *λ*_1_ = *λ*_ax_ and *λ*_2_ = *λ*_cir_. Indeed, in a biaxial extension context, we use the stretch as a direction diagonal component of the deformation gradient **F**, where *λ*_1_ = **F**(1, 1) = *F*_11_ represents the axial direction, and *λ*_2_ = **F**(2, 2) = *F*_22_ represents the circumferential direction. We assume that the tissue is perfectly incompressible, i.e., 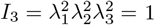, leading to **F** = diag {*λ*_1_, *λ*_2_, (*λ*_1_*λ*_2_)^−1^}. We can then rewrite equations 2 as

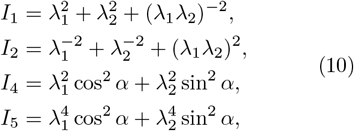

and their derivatives using equations (7) as

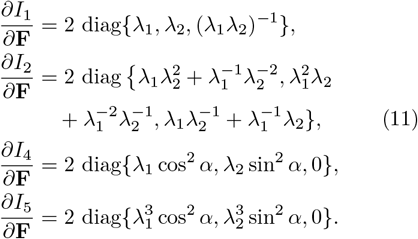

We can now insert equations (10) and (11) into the general invariant-based definition of the engineering stress (6). By assuming a zero engineering stress condition throughout the thickness, we can additionally say that *P*_33_ = 0 or **P** = diag {*P*_11_, *P*_22_, 0}. The latter defines the boundary condition for the incompressibility requirement and leads to the following expression for the hydrostatic pressure term *p*:

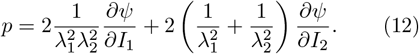

Altogether, we can explicitly provide an analytical expression for the engineering stresses *P*_11_ and *P*_22_, in terms of the applied biaxial stretches *λ*_1_ and *λ*_2_, the stretches in the axial and circumferential directions,

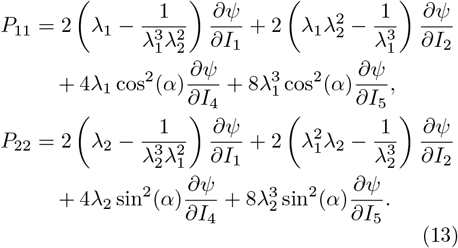

We implement two symmetrical collagen fiber families with respect to the circumferential direction, an assumption omnipresent in biomechanical analyses (Holzapfel, 2008). The latter has also been confirmed by microstructural analyses, demonstrating a symmetrical probability distribution of collagen fibers around the circumferential axis in ovine main pulmonary arteries (Rolf-Pissarczyk and Terzano, 2023). However, a clear and unique collagen fiber angle direction remains highly uncertain. Aim of this manuscript is therefore also to computationally analyse this distribution through a sweep collagen fiber angle possibilities, in order to discover the most versatile constitutive model, and further informing the directional spread. All of this without unnecessarily increasing the parameter space with multiple symmetrical fiber families. We therefore also assume the two symmetrical collagen fibers to have identical mechanical properties. We can then combine their effects in the fourth- and fifth-invariants, by multiplying the *I*_4_ and *I*_5_ contributions in equations (13) by a factor of two.

### 2.4 Model discovery

We now explore the training of our constitutive artificial neural network in Fig. 3 using different learning schemes. In particular, we will consider single or multiple samples in the loss function, with or without L1 lasso regularization. Moreover, we will compare fixing or fitting the fiber angle *α*, influencing the contribution of the anisotropic terms *I*_4_ and *I*_5_. We will consistently train all networks for 8,000 epochs, with a batch size of 32. We allow for early stopping within 2,000 epochs of no accuracy change. The robust adaptive Adam algorithm is used for first-order optimization. To prevent local minima effects, we initialized the weights of the neural network layers with random values drawn from a uniform distribution. More specifically, The Glorot normal initializer is used for the identity functions, and an unseeded random uniform initialization is used for the exponential functions, with a minimum and maximum weight value of 0.0001 and 0.1 respectively. Finally, we assess the results in terms of the individual R^2^ values in the axial and circumferential directions, for each applied ratio.

#### 2.4.1 Sample-specific model discovery

With our set of weights 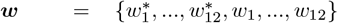 from equation (9), we perform a gradient descent learning on a weighted least-squared error loss function *L*, penalizing the error between the discovered model **P**(**F**_*i*_, ***w***) and the data 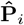:

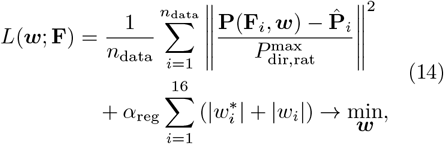

where *n*_data_ is the total number of data points of a specific sample, considering the three axial-vscircumferential experimental loading ratios, rat = {2 : 1, 1 : 1, 1 : 2} and the two axial and circumferential loading directions, dir = {ax, circ}. For our sample-specific neural networks, the number of discrete data points *n*_data_ thus corresponds to the single sample experimental results of Fig. 2, i.e., the preconditioned axial and circumferential curves of the last loading cycle for each of the three loading ratios. To account for all experiments equally, we weight the error with the maximum engineering stress per ratio and per direction,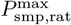.

We enforce the weights to always remain nonnegative, ***w***≥ 0. L1 regularization, or lasso regularization, enables extra feature selection and induces sparsity by reducing some weights exactly to zero, which effectively reduces model complexity and improves interpretability (McCulloch et al, 2024). Here, we set the parameter *α*_reg_ to 0.01 when L1 regularization is activated.

#### 2.4.2 Cross-sample feature selection

To include all *n* = 8 samples during training, we adapt the loss function as follows,

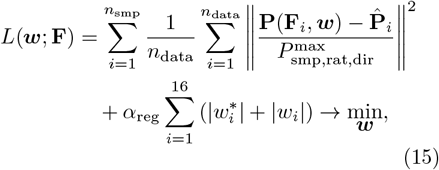

where we extend the loss function with *n*_smp_ to account for different samples, from different biaxial datasets. Instead of the extra summation, we could alternatively also use *n*_all_ instead of *n*_data_ equation (14), with *n*_all_ now equal to the total number of data points of all samples combined, for all ratios and in each direction. Note that we additionally calculate the maximum engineering stress 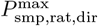 for each sample, still per ratio and per direction. This updated loss function (15) allows us to discover a unique model or set of invariant features, while we can still fit sample-specific weights in a second iteration. Through cross-sample feature selection regularization, this one-size-fits-all approach can therefore find a universal constitutive behavior. We set the parameter *α*_reg_ to 0.001 when L1 regularization is activated.

#### 2.4.3 Traditional model fitting

The traditional Holzapfel-Gasser-Ogden (HGO) model for arterial tissue assumes strain-stiffening fibers embedded in an isotropic matrix (Holzapfel et al, 2000). Considering the invariants, 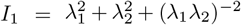 and 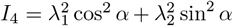, the HGO model combines the isotropic linear neo-Hookean term of the first-invariant [*I*_1_ − 3] with an anisotropic quadratic exponential term of the fourth-invariant exp([*I*_4_ − 1]^2^). The specific Holzapfel-Gasser-Ogden strain energy density function is then equal to

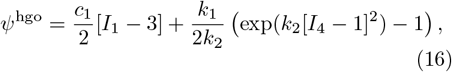

with the shear modulus *c*_1_, the stiffness-like parameter *k*_1_ *>* 0 and the non-dimensional coefficient *k*_2_ *>* 0. The last parameter highlights the exponential stiffening of the collagen fibers. Again, these three material parameters are enriched by the structural fiber angle parameter *α* in the fourth-invariant term. From our constitutive artificial neural network and the generalized equation (9), we derive and learn the exact same material model with all other weights set to zero,

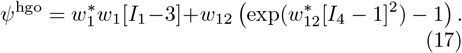

The learned weights of this model then relate oneto-one to the material parameters of the original HGO model, equation (16) with 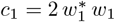 and 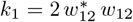 and 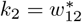.

#### 2.4.4 Reduced model discovery

After training the networks by minimizing the loss functions (14) and (15), we can reduce the complete strain energy density function of equation (9) to a select group of constitutive features or number of terms. Interestingly, we uniformly discover a two-term model for main pulmonary artery tissue with an isotropic exponential first-invariant term, (exp([*I*_1_ − 3] − 1)), and an anisotropic quadratic fifth-invariant anisotropic term, ([*I*_5_− 1]^2^). By constraining all other weights to zero during training, we can use our constitutive neural network to perform a parameter identification for the following reduced strain energy density function,

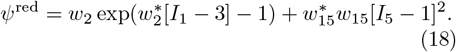

Per sample, or for all samples together, we can now fit two distinct sets of weights, 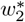 and *w*_2_ for the isotropic term, exp([*I*_1_ − 3] − 1), and 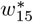 and *w*_15_ for the anisotropic term, [*I*_5_ − 1]^2^.

#### 2.4.5 Fiber angle direction

We initialize the main collagen fiber angle to a certain direction in the axial-circumferential plane and then considered it either as fixed or as fitted during network training. The parameter *α* is mathematically present in the undeformed collagen fiber vector ***n***_0_ = [cos *α*, sin *α*, 0]^*T*^, and microstructurally visible in Fig. 1. Subsequently, this microstructural information propagates into the neural network through the anisotropic *I*_4_ and *I*_5_ invariants, and appears as cos^2^(*α*) for the axial direction and sin^2^(*α*) for the circumferential direction in the engineering stress calculations of equations (13). To initialize the fiber orientation, we rely on previous microstructural analyses for biologically inspired collagen fiber directions (Fata et al, 2013; Rolf-Pissarczyk and Terzano, 2023). Researchers have used multiphoton microscopy and second-harmonic generation imaging on similar main pulmonary arteries to measure probability density functions for the collagen fiber orientation. Although these probability density functions exhibit notable in-plane axial-circumferential spreads, we postulate that the collagen fibers have a predominantly circumferential orientation.

For a fixed fiber direction, we can also vary *α* stepwise, from a complete axial orientation of *α* = 0° towards a complete circumferential orientation of *α* = 90° as {0°, 10°, 20°, 30°, 40°, 45°, 50°, 60°, 70°, 80°, 90°}. We note that for *α* = 45°, the fiber contributions to the axial and circumferential directions are identical. In a biaxial tensile test, such a scenario would lead to similar force readings in two directions, and as such a material that can be considered to showcase perfectly isotropic behavior. For the fitted fiber direction, we consider the same range of angles as a multi-start initialization or initial guess, but then learn the unknown fiber direction during training.

## 3 Results and Discussion

We report the outcomes of the different learning schemes for our constitutive artificial neural network (Fig. 3). In particular, we considered a fixed or fitted the collagen fiber angle during training, and with or without L1 regularization in the loss function. We trained the model per sample individually (equation (14), but also for all biaxial samples together (equation (15). A detailed overview of all discovered models and corresponding parameters, for all *n* = 8 samples and for all training conditions, can be found in the Supplementary Materials.

### 3.1 Sample-specific model discovery

In this section, we illustrate the process of model discovery for one representative sample, corresponding to the sample dataset in Fig. 2. We distinguish four learning variations or categories:

- training with no regularization: *α*_reg_ = 0, and fitted fiber angle *α*
- training with L1 regularization: *α*_reg_ *>* 0, and fitted fiber angle *α*
- training with no regularization: *α*_reg_ = 0, and fixed fiber angle *α*
- training with L1 regularization: *α*_reg_ *>* 0, and fixed fiber angle *α*

Training the neural networks in this section took about 4 minutes on a standard laptop for a single sample, with one condition of regularization and one initial guess of fiber angle, either fitted or fixed during training. Inspired by microstructural analyses, we plot the results for initial collagen fiber direction *α* = 70°, but subsequently train the network with a fixed or fitted angle (Rolf-Pissarczyk and Terzano, 2023).

Fig. 4 shows the discovered models and resulting engineering stress contributions of the trained weights for our selected sample, without regularization and with a fitted fiber angle. We discover a model in three terms, exp([*I*_1_ − 3]) and exp([*I*_5_ − 1]) and [*I*_5_ − 1]^2^, with an overall goodness of fit of *R*^2^ *>* 0.99. The resulting fiber angle direction is equal to 49.8° after training. The latter reflects an almost perfectly isotropic behavior, slightly oriented towards the circumferential direction. Even without any regularization, the majority of weights already train to zero. This suggests that the architecture of our neural network in Fig. 3 naturally supports sparse solutions, likely because of its orthogonal activation functions.

**Fig. 4.**
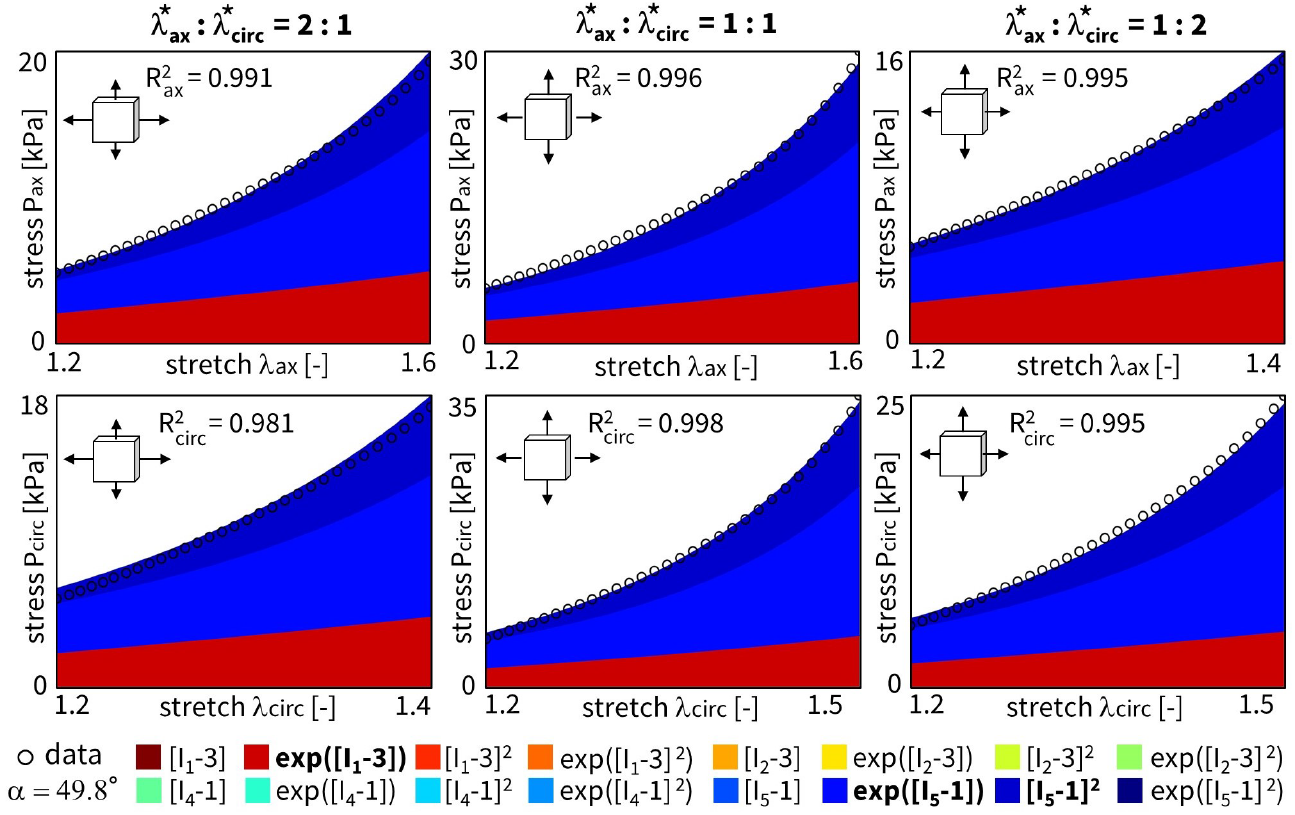
No regularization fitted fiber angle. Automated model discovery for a single sample using the full network without regularization (*α*_reg_ = 0). The legend highlights the discovered invariant terms in bold. The collagen fiber angle was *fitted* during training: 49.8°. 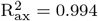 and 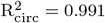.

When adding regularization to the exact same network through the L1 term (*α*_reg_ = 0.01), but still allowing the collagen fiber angle to be free and included in the learning process, we observe the results in Fig. 5. In agreement with our intuition, now, even more weights train to zero until, in the extreme case, only a single term, [*I*_5_ − 1]^2^, remains. This model reduction comes at the price of a major drop in the goodness of fit, R^2^ *<* 0.84. In support of our unregularized solution, the fiber angle remains at *α* = 49.0°, indicating a nearly isotropic behavior with a slight preference towards the circumferential direction. From a biological perspective, a one-term model in terms of only the fifth-invariant is non-intuitive, especially considering the presence of both isotropic elastin and anisotropic collagen throughout the arterial wall. These observations suggest that, with the chosen regularization parameter *α*_reg_, we have likely over-regularized the model discovery.

**Fig. 5.**
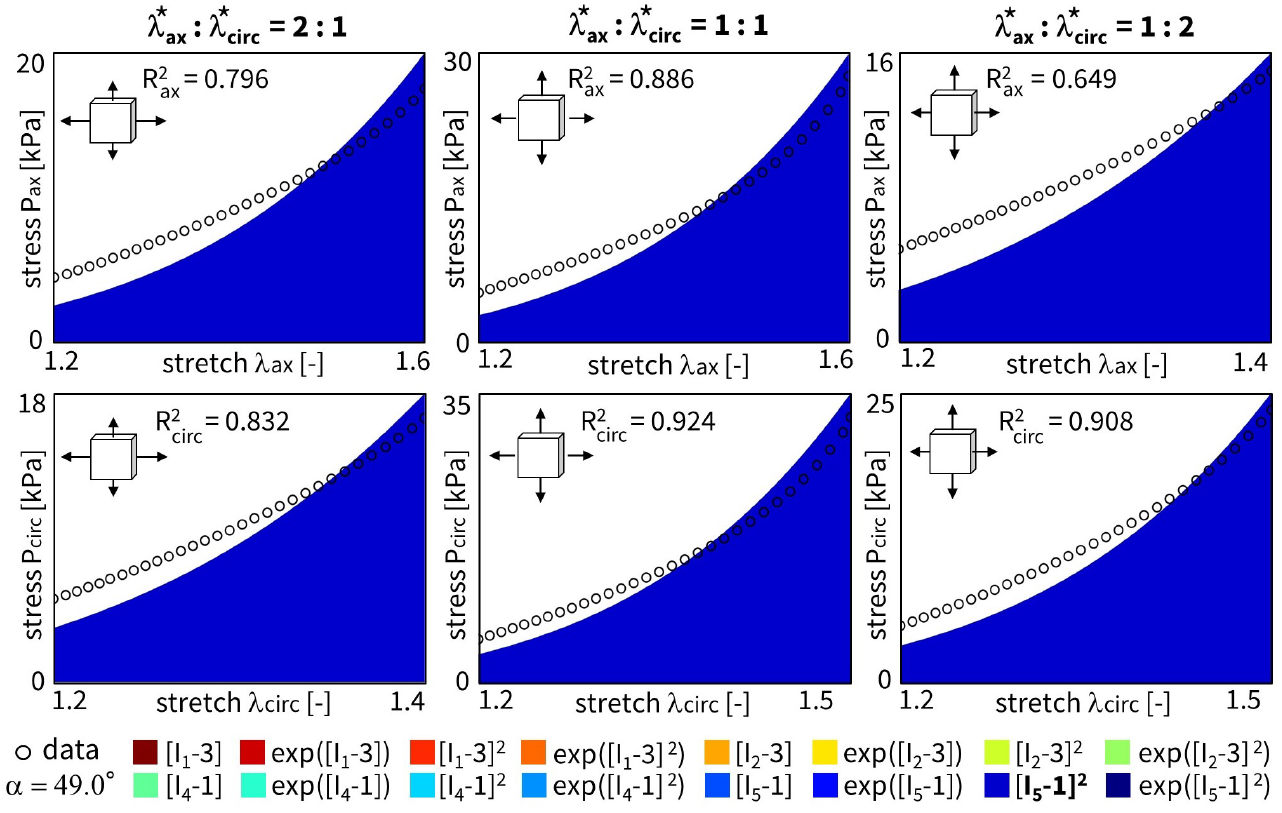
L1 regularization - fitted fiber angle. Automated model discovery for a single sample using the full network, with L1 regularization (*α*_reg_ = 0.01). The legend highlights the discovered invariant terms in bold. The collagen fiber angle was *fitted* during training: 49.0°. 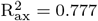and 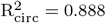.

Fig. 6 shows the results for the same input data, but now with the collagen fiber angle fixed at 70° during training. We observe a good fit with *R*^2^ *>* 0.98, and a clear, more substantial contribution of the isotropic terms and weights. Specifically, we discover the full range of first-invariant terms, [*I*_1_ − 3] and exp([*I*_1_ − 3]) and [*I*_1_ − 3]^2^ and exp([*I*_1_ − 3]^2^), while the anisotropic terms [*I*_5_ − 1] and [*I*_5_ − 1]^2^ now only have minor contributions.

**Fig. 6.**
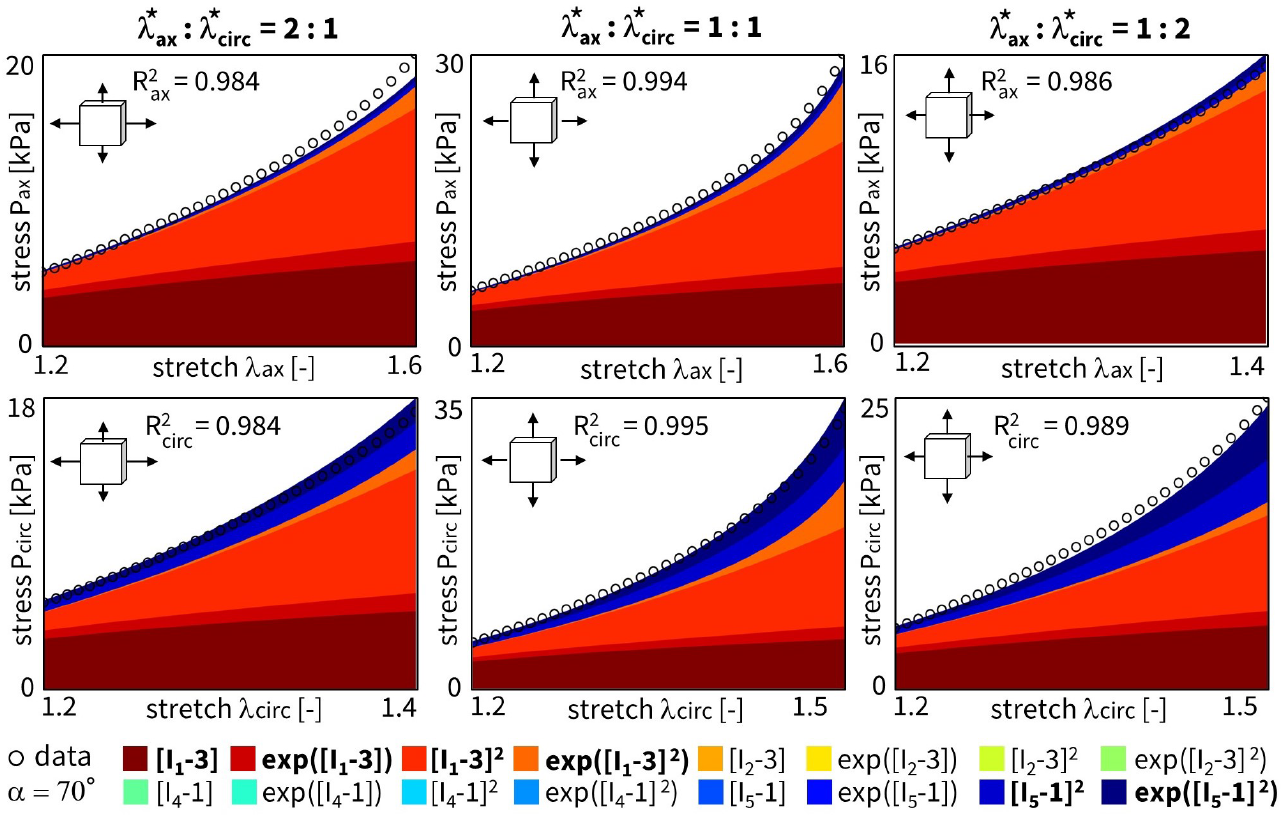
No regularization - fixed angle. Automated model discovery for a single sample using a full network without regularization (*α*_reg_ = 0). The legend highlights the discovered invariant terms in bold. The collagen fiber angle was circumferentially *fixed* during training (70°). 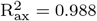and 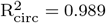.

We can again regularize the same network using L1 regularization ((*α*_reg_ = 0.01), while fixing the collagen fiber angle *α* at 70°. Fig. 7 clearly shows the expected reduction in the number of discovered terms. The isotropic exponential term, exp([*I*_1_ − 3]), and the anisotropic quadratic term, [*I*_5_ − 1]^2^, which emphasizes the circumferential direction, are now more evident. By comparing the regularized network with fixed angle in Fig. 7 with the regularized network with fitted angle in

**Fig. 7.**
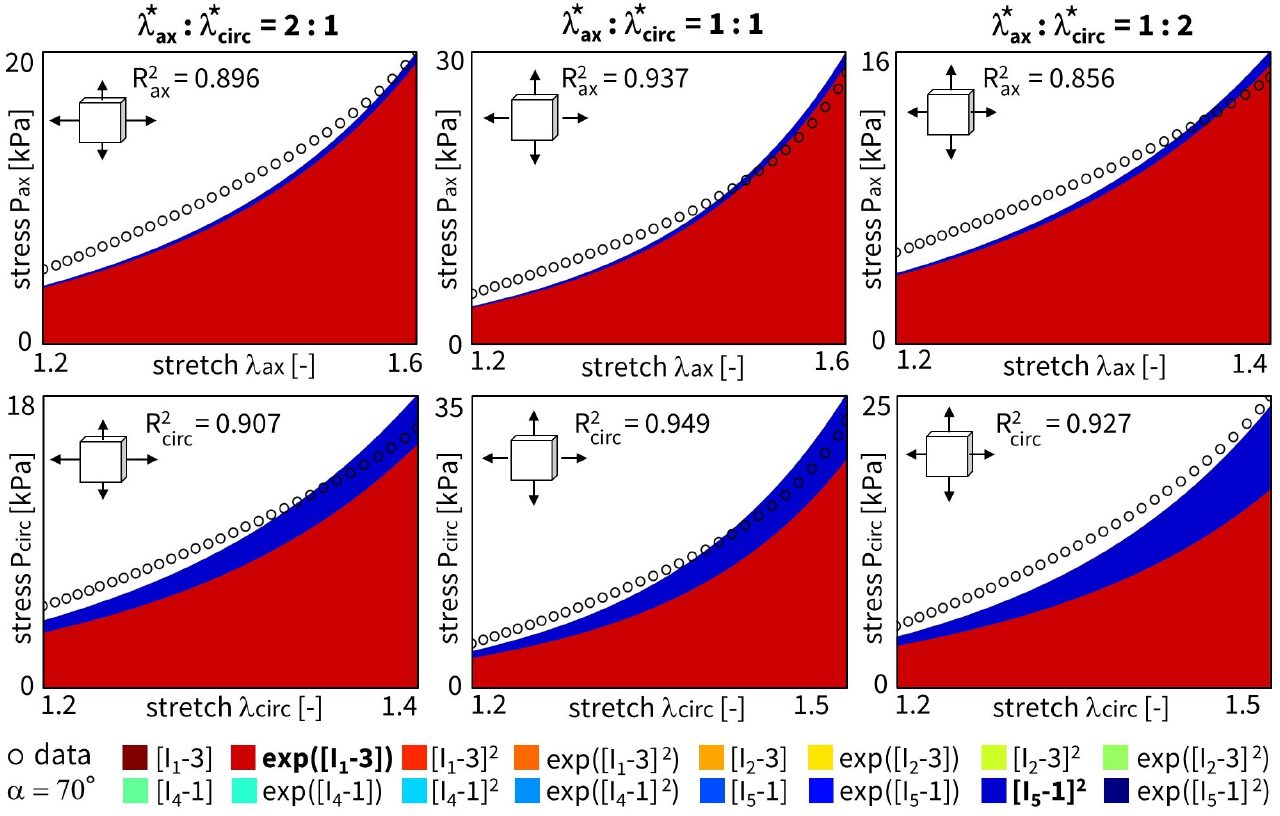
L1 regularization - fixed angle. Automated model discovery for a single sample using a full network with L1 regularization (*α*_reg_ = 0.01). The legend highlights the discovered terms in bold. The collagen fiber angle was circumferentially *fixed* during training (70°). 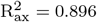 and 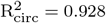.

Fig. 5, we can clearly see a major isotropic *I*_1_ presence and a minor circumferential anisotropic *I*_5_ contribution when the fiber angle is fixed. When the angle direction is fitted, the model adapts towards a more isotropic angle, with larger, but microstructurally less relevant *I*_5_ weights.

### 3.2 Cross-sample feature selection

Following section 2.4.2 on the regularization through cross-sample feature selection, and according to the updated loss function over all samples of equation (15), we train a network with all *n* = 8 samples combined. The latter corresponds to locations A and B in Fig. 1, originating from the four distinct ovine pulmonary roots. Naturally, all samples are now taken into account. To reduce uncertainty across the samples, we fix the fiber angle to 70°. Here, our objective is to discover one single model for all data sets. This implies that the magnitude of the weights for the individual samples is of secondary importance. For all samples combined, we can now plot the results without regularization (Fig. 8) and with L1 regularization (Fig. 9).

**Fig. 8.**
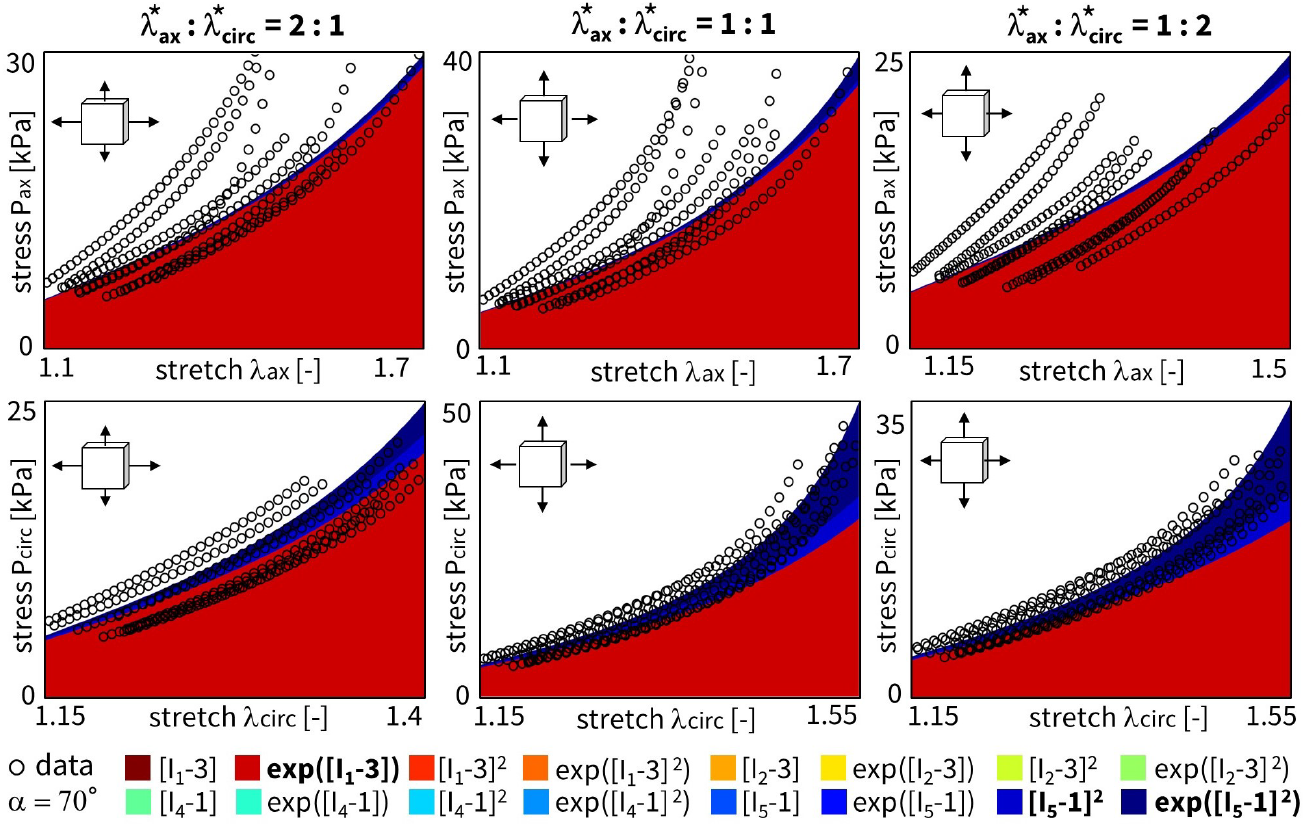
Cross-sample feature selection – L1 regularization. Automated model discovery for all samples combined, without L1 regularization (*α*_reg_ = 0). The collagen fiber angle was circumferentially *fixed* during training (70°). The legend highlights the non-zero weights in bold.

**Fig. 9.**
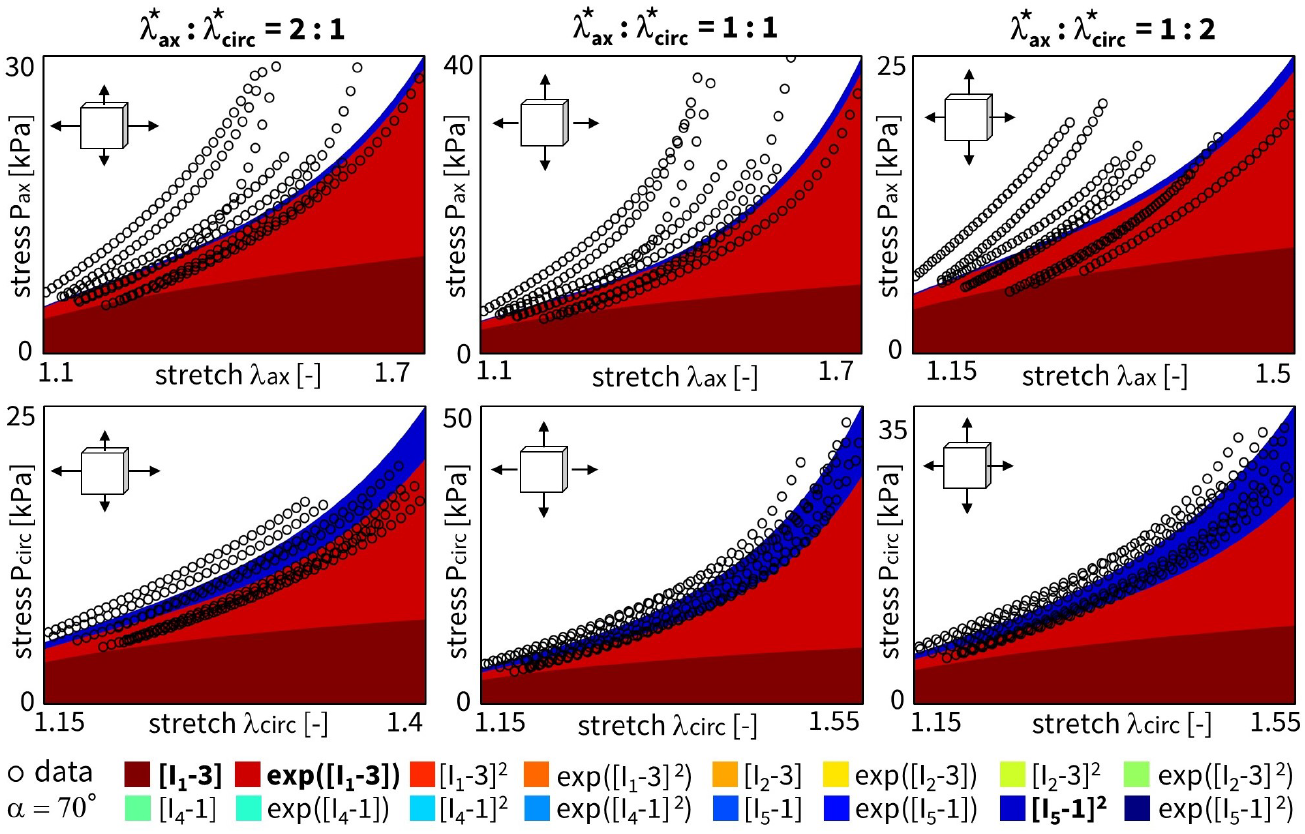
Cross-sample feature selection – L1 regularization. Automated model discovery for all samples combined, with L1 regularization (*α*_reg_ = 0.001). The collagen fiber angle was circumferentially *fixed* during training (70°). The legend highlights the non-zero weights in bold.

The discovered features are as follows:

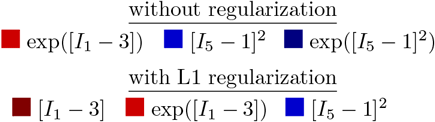

Here, we have lowered the impact of the L1 regularization by reducing the regularization parameter to *α*_reg_ = 0.001. Notably, we consistently discover an isotropic and an anisotropic term to describe all eight samples combined. Without regularization, and out of more than 60,000 models, we discover the exact same constitutive behavior as previous studies for layers of healthy human aortas (Peirlinck et al, 2024b). Indeed, for an experimental dataset originating from tissues under different native blood pressure conditions (pulmonary vs. aortic tissue), the neural network overcomes aleatoric and epistemic uncertainties to discovery the exact same constitutive behavior. Taking both cross-sample feature selection training schemes, we observe an overlap of the same exp([*I*_1_ −3]) and [*I*_5_ −1]^2^ features, without and with regularization, respectively. Training this enlarged neural network for all *n* = 8 samples takes about 20 minutes per condition of regularization. As part of the network’s design, we initially train the weights in a one-size-fits-all approach. This will be useful for representing the entire population of main pulmonary arteries, and will allow us to retrain the weights while constraining the network to the universally discovered feature invariants. The Supplementary Materials additionally present the cross-sample feature selection results for different initial fiber angle directions, both fixed or fitted during training.

### 3.3 Fiber angle analysis

Considering all individual *n* = 8 samples, we now train the network for a sweep of fixed fiber angles. As such, we can perform an in-depth sensitivity analysis on this collagen fiber angle, sweeping the entire axial-circumferential domain. Fig. 10 shows the results without L1 regularization, highlighting the predominantly discovered isotropic exponential term, exp([*I*_1_ − 3]). Clearly, a higher number of anisotropic terms are discovered when the fiber angle is fixed toward a more circumferential range of the sweep with *α >* 45°. The latter corresponds to the observed increased non-linearity in the circumferential direction of our data. In other words, we observe a naturally occurring sparsity in the anisotropic invariants for the more compliant direction, while training in the stiffer direction picks up the anisotropic invariant terms. Fig. 11 shows the same training workflow across angles but now with L1 regularization applied. When comparing both histograms, we observe a clear sparsification of our regularization through a decrease in the number of discovered terms, along with a shift from isotropic to anisotropic terms. The anisotropic quadratic fifth-invariant term, [*I*_5_ − 1]^2^, becomes the most prevalent now. Again, we see an increased sparsity in the anisotropic terms for the axial range of fiber angles for *α <* 45°. The main feature discovery when sweeping the collagen fiber angle can be summarized as follows:

**Fig. 10.**
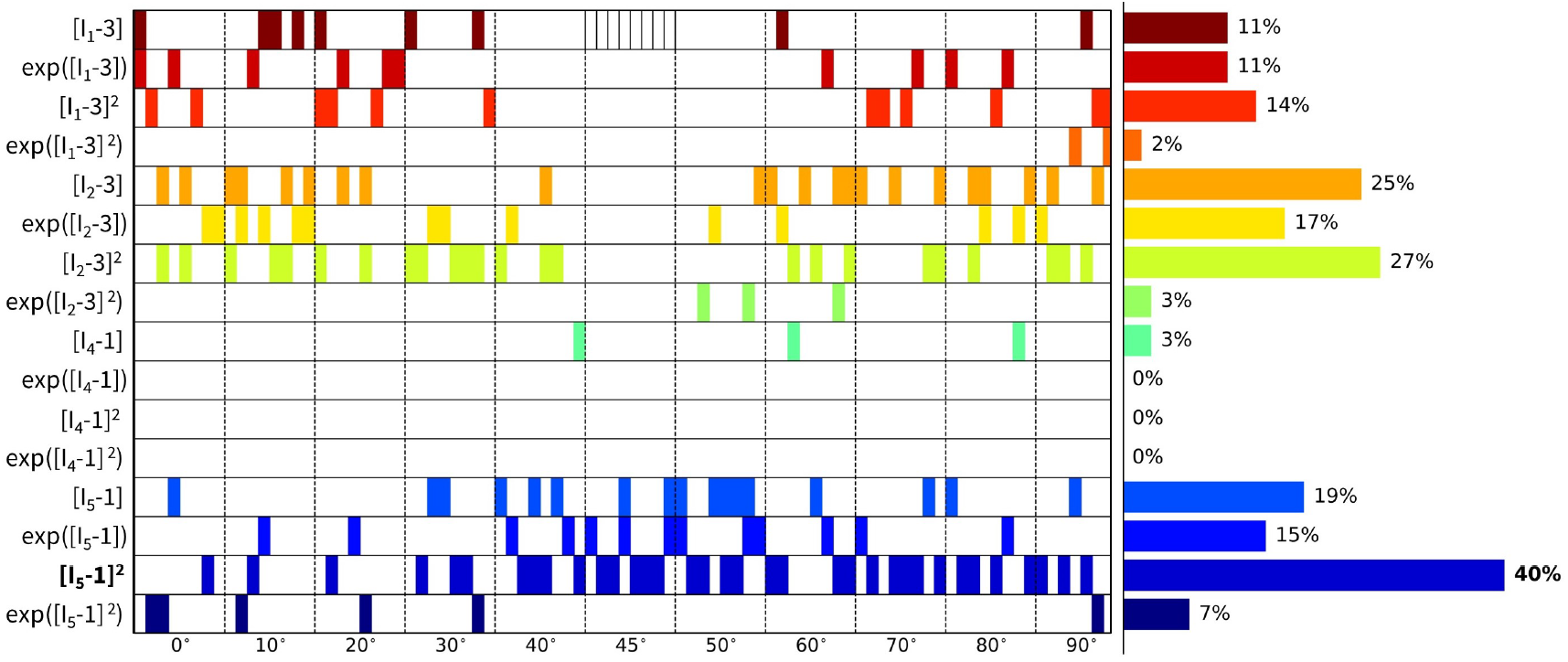
Fixed fiber angle sweep - no regularization. Sweep of fixed collagen fiber angles, without regularization (*α*_reg_ = 0). We vary the collagen angle in the following 11 steps: {0°, 10°, 20°, 30°, 40°, 45°, 50°, 60°, 70°, 80°, 90°}, where 0° represents the axial fiber direction, 45° indicates isotropic fiber behavior, and 90° corresponds to a perfect circumferential fiber direction. A separate network for each of the *n* = 8 samples has been trained in each collagen fiber angle category. The legend highlights the discovered terms and their prevalence across the whole 11 angles x 8 samples = 96 training routines spectrum, predominated by exp([*I*_1_ − 3]).

**Fig. 11.**
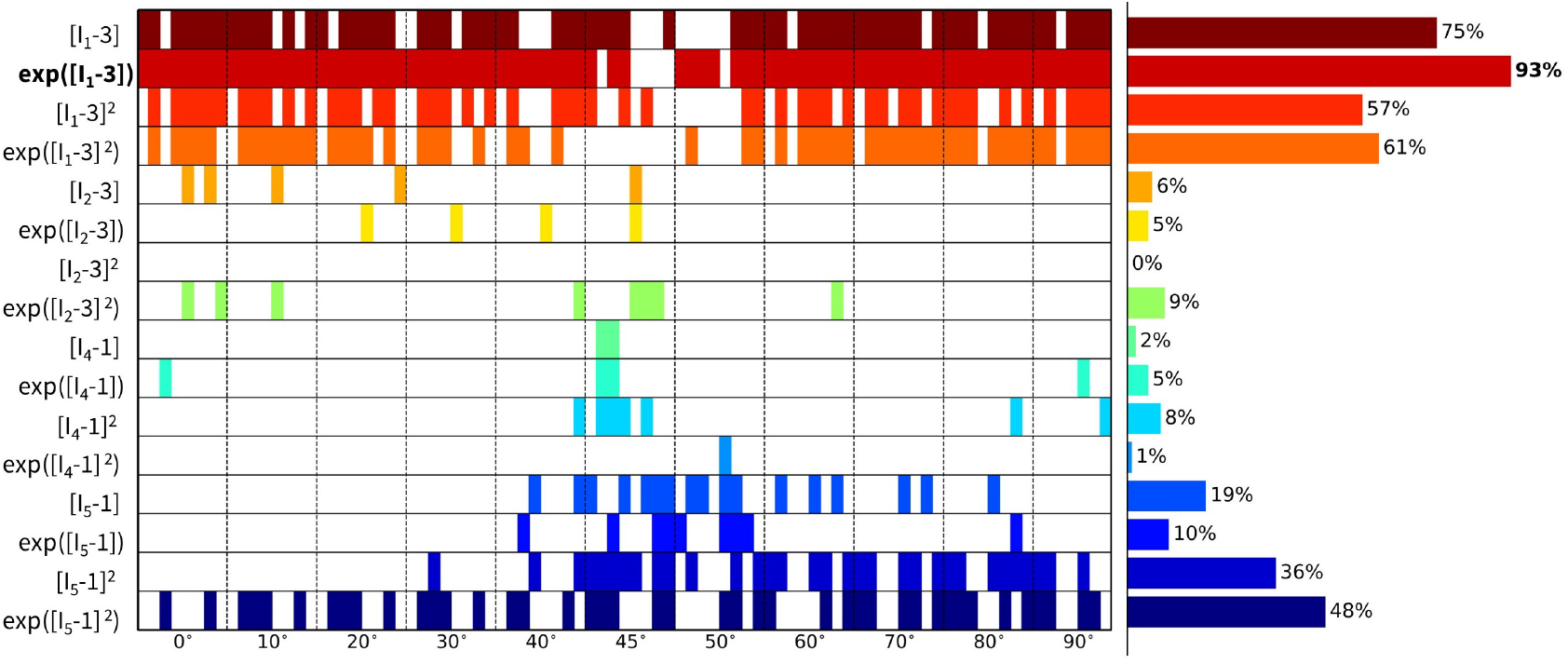
Fixed fiber angle sweep - L1 regularization. Sweep of fixed collagen fiber angles with L1 regularization-(*α*_reg_ = 0.01). We vary the collagen angle in the following 11 steps: {0°, 10°, 20°, 30°, 40°, 45°, 50°, 60°, 70°, 80°, 90°}, where 0° represents the axial fiber direction, 45° indicates isotropic fiber behavior, and 90° corresponds to a perfect circumferential fiber direction. A separate network for each of the *n* = 8 samples has been trained in each collagen fiber angle category. The visual classification of the different samples per angle group is indicated in the cell of the first row, middle column. The legend highlights the discovered terms and their prevalence across the whole 11 angles x 8 samples = 96 training routines spectrum, predominated by [*I*_5_ − 1]^2^.

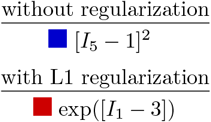

Interestingly, we also observe a general shift from the isotropic first-invariant *I*_1_ terms towards the more nonlinear isotropic second-invariant *I*_2_ terms. Recent research has also shown that the second-invariant in the isotropic part of the strain–energy function can enrich the model discovery process (Kuhl and Goriely, 2024). Moreover, this is in agreement with similar neural network methods showing that cardiac tissue is best described by second-invariant models (Martonova et al, 2024).

The dispersed Gasser-Ogden-Holzapfel model proposes a collagen fiber dispersion to overcome the assumption of single-direction fiber families (Gasser et al, 2006):

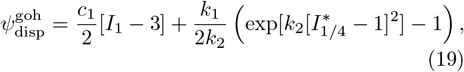

where a mixed isotropic-anisotropic fiber invariant term expands as 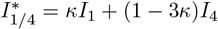,and the microstructural fiber dispersion parameter *κ* significantly influences the constitutive equation (Peirlinck et al, 2024a). When *κ* = 0, we have 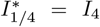,and the model reduces to the traditional HGO model in equation (17), with no collagen fiber dispersion. When *κ* = 1*/*3, we have 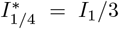,and the model reduces to a fully isotropic exponential model. A distribution of collagen fiber directions would be in agreement with the probability density function of angle directions around the circumferential axis in ovine main pulmonary arteries, as observed by Rolf-Pissarczyk and Terzano (2023). However,the focus of this work is to discover a selected set of invariants with a clear distinction between isotropic and anisotropic terms, reason why we did not include a phenomenological fiber dispersion parameter *κ*. Alternatively, we could also introduce multiple fiber families, each with different discrete fiber angles *α*_*i*_ and/or mechanical properties (Ramachandra and Humphrey, 2019). These collagen fiber families could then be incorporated into the CANNs through multiple invariants *I*_4(*ii*)_ and *I*_5(*ii*)_ (Peirlinck et al, 2024a). This approach can be implemented in the HGO model through a fiber-specific term *E*_*i*_,

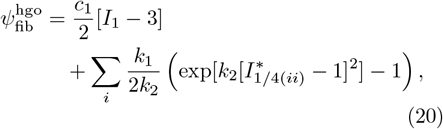

either with or without dispersion (Sempertegui and Avril, 2024).

Altogether we postulate that the collagen fibers have a predominant circumferential orientation, which also corresponds to the mechanical data, and the discovered constitutive behavior during the alpha sweeps. Indeed, anisotropic invariants become apparent only in the circumferential range of collagen fiber angles. When circumferentially fixing the collagen fiber orientation on the other hand, circumferential fibers again align with the distribution of Rolf-Pissarczyk and Terzano (2023).

### 3.4 Traditional model fitting

As a cross-comparison with literature, we now constrain our network to the traditional HGO model of equation (17), consisting of an isotropic linear first-invariant term, [*I*_1_ − 3], and an anisotropic quadratic exponential fourth-invariant term, exp([*I*_4_ − 1]^2^) (Holzapfel et al, 2000). Fig. 12 shows the resulting weights after training, with a fitted collagen fiber angle equal to 49.0°. L1 regularization is not applicable here, and we learn the fiber angle after initialization. Through equation (16), we report the known HGO model parameters based on the learned weights, *c*_1_ = 1.95 kPa and *k*_1_ = 0.77 kPa and *k*_2_ = 0.145. These parameters fall within expected stiffness ranges, and the fit can be considered fairly good. This is reasonable, as the HGO model has been widely validated for a broad range of biological tissues.

**Fig. 12.**
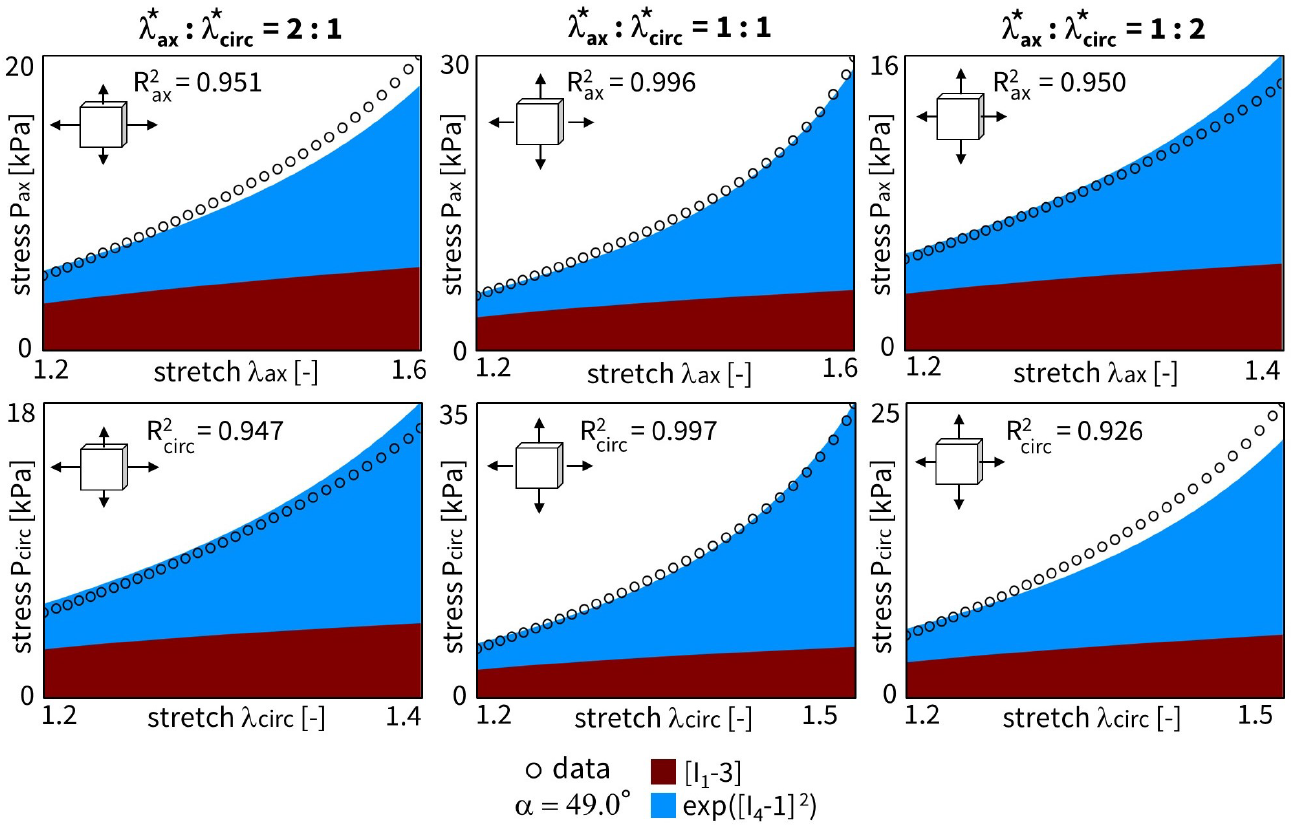
Traditional Holzapfel-Gasser-Ogden fit. Material fitting for a single sample using the HGO neural network. The *fitted* collagen fiber angle is equal to 49.0° after training. 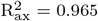 and 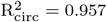. The learned weights are *w*_1,1_ = 0.838, *w*_2,1_ = 2.33 kPa, *w*_1,12_ = 0.413, and *w*_2,12_ = 2.66 kPa.

### 3.5 Reduced model discovery

Considering the results from the cross-sample feature selection and the fiber angle analysis in the previous Sections 3.2 and 3.3, we alternatively constrain our neural network from Fig. 3 or general equation (9) to only the two newly discovered terms, exp([*I*_1_ −3]) and [*I*_5_ −1]^2^. The latter would correspond to a material fitting with the reduced constitutive equation (18). The trained weights of this novel constitutive model yield a balanced isotropic-anisotropic contribution, and a general goodness of fit of *R*_2_ *>* 0.99, even with only two terms. The results for our representative sample are shown in Fig. 13. Obviously, network L1 regularization is obsolete here. Training this reduced neural network takes about 90 seconds. Note that the collagen fiber angle fitted in the current reduced model plots. As expected, we observe a slightly circumferential orientation of the fiber angle for the fifth-invariant term, but with no real pronounced anisotropy in the overall model (*α* = 50.9°). Additionally, the mechanical data themselves display a more compliant behavior in the axial direction, with higher stretches after preload correction, and a stiffer behavior in the circumferential direction, with higher experimental forces, all in agreement with the discovered models here, and with previously published experimental data on similar tissue types.

**Fig. 13.**
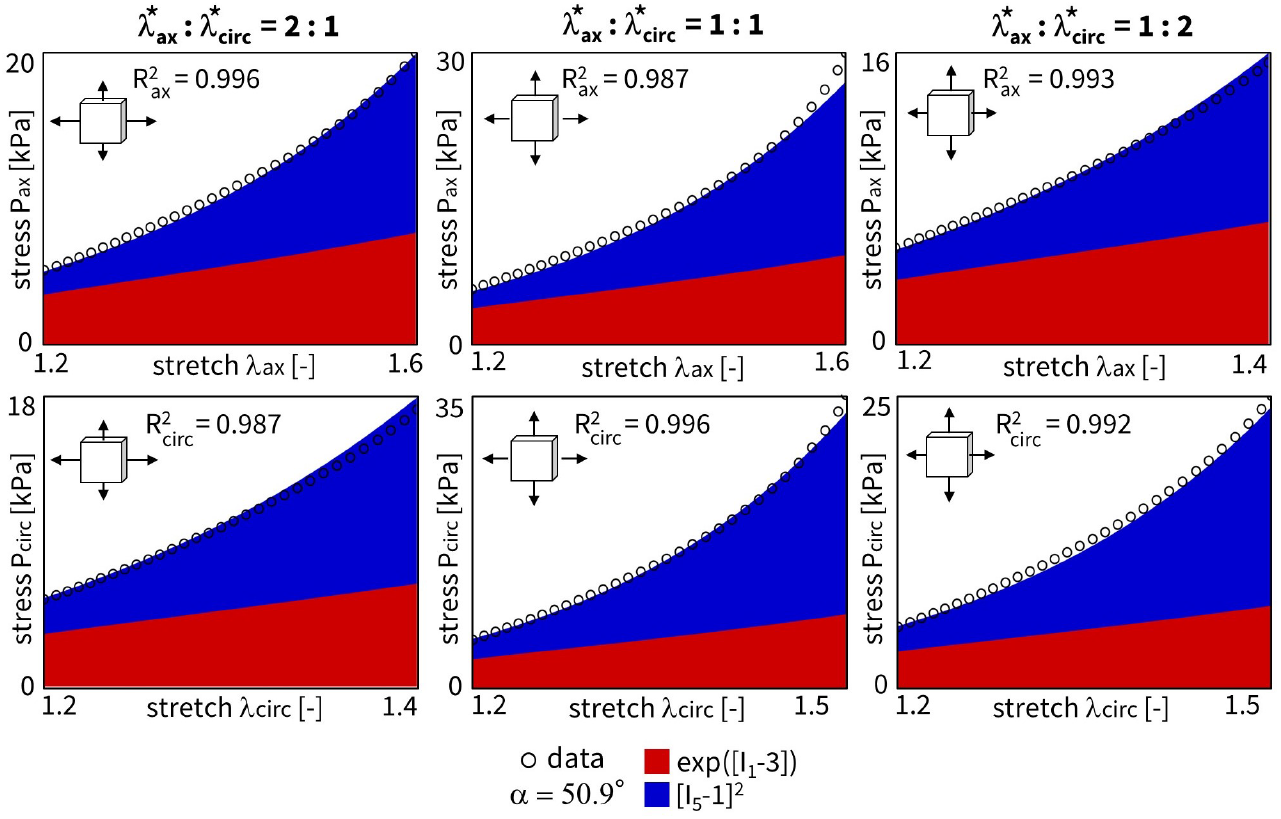
Reduced constitutive neural network fit. Material fitting for a single sample using the reduced neural network. The *fitted* collagen fiber angle is equal to 50.9° after training. 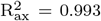 and 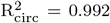. The learned weights are *w*_1,2_ = 0.185, *w*_2,2_ = 10.45 kPa, *w*_1,15_ = 0.462, and *w*_2,15_ = 0.274 kPa.

Notably, our newly discovered constitutive equation (18) leads to an increase in R^2^ compared to traditional HGO model (comparing fitting accuracy of Fig. 13 with Fig. 12). It is also striking that we rarely discover the characteristic exponential quadratic fourth-invariant term of this traditional material model when sweeping the collagen fiber angles for all samples, as shown in

Fig. 10 and Fig. 11, with a 1% and 0% occurrence without and with L1 regularization, respectively. Instead, the anisotropic quadratic fifth-invariant term of the updated reduced model seems to better minimize the loss function of our network, and is therefore better suited to characterize the generally compliant and highly non-linear constitutive behavior of pulmonary arteries. Accordingly, also an exponential first-invariant term exp([*I*_1_ − 3]) seems to be the preferred learned isotropic part of the strain–energy function, instead of the traditional linear [*I*_1_ − 3].

To further quantify our results, Fig. 14 illustrates the averaged R^2^ value for all of the *n* = 8 samples, with individual weights trained for the whole range of fixed collagen fiber angles. Heat map colors compare the fitting accuracy of the reduced model (exp([*I*_1_ −3]) and [*I*_5_ −1]^2^) to the HGO model ([*I*_1_ −3] and exp([*I*_4_ −1]^2^)). For a fiber angle in the range 45° ≤*α* ≤50°, and fixed in this direction during learning, the traditional model fits the dataset well. However, we see a clear fitting improvement and model generalization for all fiber angles with our newly discovered constitutive equation. By combining an isotropic exponential first-invariant term with an anisotropic quadratic fifth-invariant term, we observe an R^2^ *>* 0.9 over the whole collagen fiber spectrum, whereas the traditional model only performs well when the fixed angles are in the same range as the resulting fitted fiber angles (close to isotropy, tending towards the circumferential direction). The exact weights for all angles and for both models are provided in the Supplementary Materials. Both the HGO model and the newly discovered constitutive law have three distinct model parameters, with the fiber angle direction as possible extra structural variable. A unique fit is therefore unlikely, but an accurate constitutive description highly important.

**Fig. 14.**
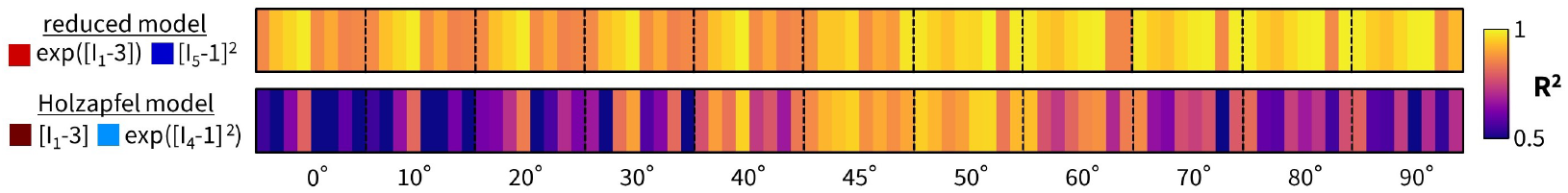
Fitting accuracy heat map comparison. Figure compares the reduced constitutive neural network fit with the traditional Holzapfel-Gasser-Ogden fit in terms of total R^2^, averaged over the axial and circumferential direction, for all ratios. The restricted neural network was trained for each of the *n* = 8 samples in each imposed fiber angle category. We vary the fixed collagen angle in following steps: {0°, 10°, 20°, 30°, 40°, 45°, 50°, 60°, 70°, 80°, 90°}, where 0° represents the axial fiber direction, 45° indicates isotropic fiber behavior, and 90° corresponds to a perfect circumferential fiber direction.

Given the uncertain probability density distribution of collagen fiber orientations observed by Rolf-Pissarczyk and Terzano (2023), it remains challenging to define a single and main collagen direction in sheep pulmonary arteries. We therefore want a constitutive model that is predictive, versatile and accurate for a range of collagen fiber angles, predominated by the circumferential direction. Our newly discovered constitutive model more has an isotropic exponential term and an anisotropic quadratic fifth-invariant anisotropic term, compared to the combination of an isotropic linear term and an anisotropic quadratic exponential fourth-invariant term in the HGO model. The former thus appears to better describe the highly non-linear behaviour of main pulmonary arteries, with preferred fiber directions symmetrically around the circumferential direction. The traditional HGO model also appears to be more sensitive to the unknown and uncertain collagen fiber angle.

Finally, we can also uniformly fit the same reduced neural network considering all samples together, as visualized in Fig. 15. We used cross-sample feature selection and training on the whole dataset to discovered a combination of an isotropic exponential first-invariant term, exp([*I*_1_ − 3]), and an anisotropic quadratic fifth-invariant term, [*I*_5_ − 1]^2^. We can subsequently test this updated constitutive model on the same complete dataset together, see equation 15, but by constraining the neural network weights accordingly now. Although a clear and unique collagen fiber angle direction around the circumferential direction remains highly uncertain, we interestingly observe a fitted fiber angle of 89.9°. Training all samples together thus results in an almost perfectly circumferential collagen fiber direction. Again, even when neglecting fiber dispersion, this is again in agreement with the experimental data Rolf-Pissarczyk and Terzano (2023).

**Fig. 15.**
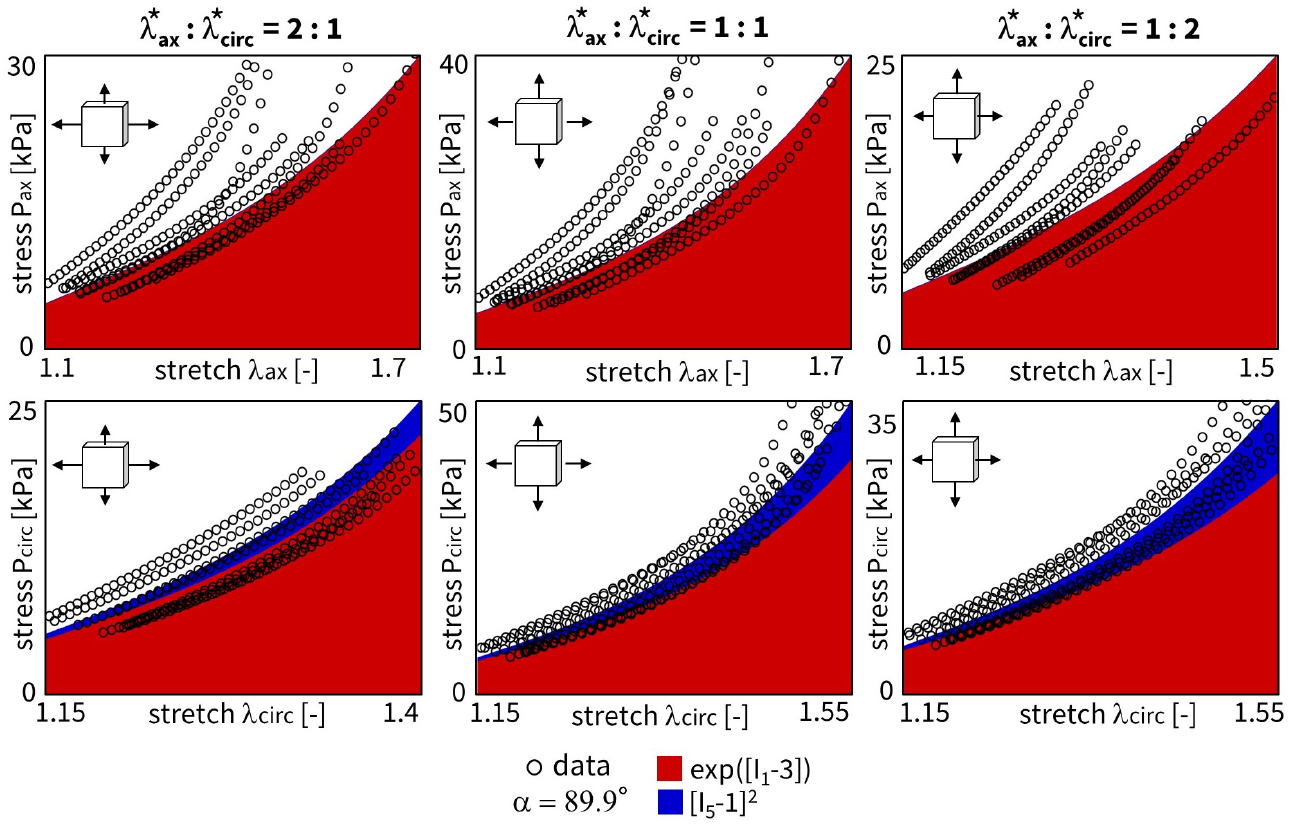
Reduced constitutive neural network all samples. Unique material fitting using the reduced neural network for all samples combined. The *fitted* collagen fiber angle is equal to 89.9° after training. The learned weights are *w*_1,2_ = 0.424, *w*_2,2_ = 8.48 kPa, *w*_1,15_ = 0.222, and *w*_2,15_ = 0.112 kPa.

Taken together, our invariant-based model discovery approach with more than 60,000 possible model terms, reduced to a select subset of constitutive behaviors, provides crucial insights into the biomechanics of the pulmonary arterial wall, that would be impossible to obtain with a traditional modeling approach.

## 4 Conclusion and outlook

In this work, we discovered a novel constitutive model for the main pulmonary arteries:

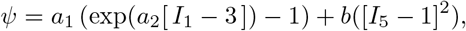

characterized by three material parameters, *a*_1_, *a*_2_ and *b*, which aligns perfectly with our recently discovered material model for systemic arterial tissues (Peirlinck et al, 2024b). Specifically, we combined biaxial tensile testing of *n* = 8 pulmonary artery samples and automated model discovery using constitutive neural networks. Our constitutive neural network approximates the free energy function *ψ* using sixteen terms. Its first layer generates powers (°) and (°)^2^ of the four network invariants, *I*_1_, *I*_2_, *I*_4_, *I*_5_, while the second layer applies the identity (°) and exponential function (exp(°)) to these powers. From cross-sample feature selection regularization and an in-depth sensitivity analysis of the collagen fiber direction, we conclude that the constitutive behavior of pulmonary arteries is best characterized by a two-term model in terms of an isotropic exponential first-invariant term and an anisotropic quadratic fifth-invariant term.

Our main pulmonary artery biaxial dataset allowed us to discover the updated constitutive material model in supra-physiological strain regimes, relevant for the Ross procedure. Computational models of the Ross procedure could now allow us to simulate the actual *in vivo* conditions, and estimate wall stresses and deformations over time. This would allow us to improve clinical outcomes by investigating and predicting potential chirurgical solutions. Key and critical for such *in silico* simulations would then again be the constitutive description of the considered pulmonary tissue. Moreover, an accurate material model is especially important in regions of high stress concentrations (Peirlinck et al, 2024b). This is to be expected in the context of the Ross procedure, where pulmonary arteries are acutely exposed aortic conditions, i.e. four to fivefold increase in blood pressures.

Future work should first address the uncertainty in the data, possible through recently developed Bayesian constitutive artificial neural networks (Linka et al, 2025). This would alleviate the averaging assumptions of the presented cross-sample feature selection regularization. Moreover, we could explicitly learn the uncertainty in the fiber direction with a mean collagen fiber angle and a standard deviation that characterizes the fiber dispersion, instead of using a purely phenomenological fiber dispersion parameter.

Linka et al (2023) used the same convolutional artificial neural network approach to autonomously discover constitutive models for skin, based on very similar biaxial tensile tests. Biaxial test ratios were divided into training and test data. While the parameter values were different for each set of training data, the set of active parameters and discovered invariants were the same across all datasets, discovering a uniform constitutive model. Here, our holistic biaxial dataset is a prerequisite to cover a large stretch area in the axial-circumferential plane, and as such describe and discover our arterial tissue the best. Future work could therefore consider a validation step through a different mechanical setup, such as combined extension-inflation test (Wang et al, 2021). Pressure-diameter curves could indeed nicely relate experimental and *in vivo* condition with respect to axial and circumferential stretches. A computational model with the newly discovered constitutive description of main pulmonary arteries could then show that we indeed better predict stresses and strains for untrained data.

Similar artificial neural networks have recently enabled the robust discovery of anisotropic constitutive models for a wide variety of warp-knitted fabrics (McCulloch and Kuhl, 2024). By considering different mounting orientations, the authors discovered interpretable anisotropic models that performed well in both training and testing. Such medical textiles, used in cardiovascular applications, can serve as external supports for the Ross procedure to prevent acute dilatation of the transplanted pulmonary tissue in the high-pressure aortic environment (Verbrugghe et al, 2013; Nappi et al, 2020; Vervenne et al, 2023). Many unknowns remain regarding the mechanocompatibility of these textiles and their interaction with the native tissue (Singh et al, 2015; Ramachandra et al, 2020). Textile stiffness should prevent short-term acute dilatation, while textile compliance should restore long-term blood pressure pulsatility. Once again, computational design solutions will critically depend on the models and parameters of the underlying pulmonary autograft tissue.

Our main take-home message is that the constitutive behavior of main pulmonary arteries can be best described by combining an isotropic exponential first-invariant term, exp([*I*_1_ − 3]), and with an anisotropic quadratic fifth-invariant term, [*I*_5_ − 1]^2^. Critical to discovering this sparse and interpretable model has been cross-sample feature selection and a thorough collagen fiber angle analysis sweep.

## Supplementary Materials

The source data, code, and results are available in KU Leuven’s research data repository: https://doi.org/10.48804/almeep.

## Acknowledgments

This work was supported by a doctoral fellowship SB1SE2123N from the Research Foundation Flanders (FWO) to Thibault Vervenne, by the research project G029819N from the FWO to Nele Famaey, by the NWO Veni Talent Award 20058 to Mathias Peirlinck, by the NSF CMMI Award 2320933 and the ERC Advanced Grant 101141626 to Ellen Kuhl. Additional travel grants from the FWO and the R. Snoeys Foundation to Thibault Vervenne made this collaboration possible.

## Declarations

